# *Ex Vivo* Delivery of mRNA to Immune Cells *Via* a Non-Endosomal Route Obviates the Need for Nucleoside Modification

**DOI:** 10.1101/2024.10.31.621287

**Authors:** Bartika Ghoshal, Debajyoti Chakraborty, Manish Nag, Raghavan Varadarajan, Siddharth Jhunjhunwala

## Abstract

Base modification and the use of lipid nanoparticles (LNPs) are thought to be essential for efficient *in vivo* delivery and expression of mRNA. However, for *ex vivo* immune cell engineering, the need for either of the two is unclear. Previous reports have suggested that nucleic acids may be efficiently delivered to immune cells *ex vivo,* through a non-endosomal delivery route, but the need for base modification has not been determined. Herein, we demonstrate that when a non-endosomal delivery method is used, unmodified mRNA performs equally well to the commonly used base-modified mRNA, including the *N*^1^ methyl pseudo-uridine modification, in terms of protein expression and inflammatory response in cells. However, if an endosomal delivery route is used, then *N*^1^ methyl pseudo-uridine modification is necessary for high expression and low inflammatory response, as demonstrated by others as well. Overall, we show that non-endosomal mRNA delivery renders nucleoside modifications non-essential, and that unmodified mRNA combined with non-endosomal delivery route may be used for efficient *ex vivo* mRNA-based engineering of immune cells.

## Introduction

Messenger Ribonucleic Acid (mRNA)-based therapeutics have emerged as a potential line of treatment for multiple diseases. While mRNA molecules are known for their instability and high immunogenicity, Kariko et al provided a breakthrough by introducing base modifications, which improved stability and reduced activation of the immune system^1^. Immune cell activation was thought to primarily occur through activation of Toll-Like Receptors (TLR) present in the endosomes of these cells^2^. To reduce the TLR-based activation, among the many possible nucleoside modifications, Ψ (pseudo uridine) in place of uridine resulted in low immunogenicity, better stability and translational capacity of the mRNA^3,4^. Further improvements demonstrated that *N*^1^ methyl Ψ works better than Ψ in terms of both stability and lower immune response of the resulting mRNA^5,6^. Hence, this modification has become the most used in mRNA therapeutics.

However, the need for extensive base modifications has been questioned more recently. For example, Kauffman et al have reported that if the therapeutic mRNA is encapsulated in a lipid nanoparticle, then the unmodified and Ψ modified mRNA show no difference in immunogenicity or expression, *in vivo*^7^. The lipid nanoparticle was thought to protect the mRNA from both serum degradation and reduce its interaction with endosomal TLRs, hence rendering the need for modification unnecessary. This led us to ask the following question: are base modifications necessary only if the mRNA is delivered through the endosomal route and by extension not required if the delivery is non-endosomal?.

To address this question, in the present study, we evaluate the immunogenicity and expression levels of unmodified or modified mRNA delivered through the endosomal route and compare them to those delivered *via* the non-endosomal route in immune cells *in vitro*. Our data show that non-endosomal delivery obviates the need for base modifications in mRNA, as both unmodified and modified mRNA show similar expression levels and minimal activation of immune responses. However, if the delivery is through the endosomal route, then base modification, especially with *N*^1^ methyl Ψ, is necessary for better expression and lower immune response. In summary, this study provides evidence for the need for mRNA modification if the delivery is through the endosomal route, but not for non-endosomal delivery. This has implications for *ex vivo* modification of immune cells through delivery of exogenous mRNA.

## Materials and Methods

### Cell culture

The mouse macrophage cell line, RAW 264.7 (RAW), was cultured in Dulbecco’s Modified Eagle’s medium (DMEM; MP Biomedicals, California) with 10% heat-inactivated fetal bovine serum (FBS-Gibco, Thermo Fisher Scientific) and 1% Antibiotic-Antimycotic (Anti-Anti-Gibco, Thermo Fisher Scientific). Human neutrophil-like cell line, HL60, was cultured in Iscove’s Modified Eagle’s medium (IMDM-Sigma Aldrich), 20% FBS and 1% Anti-Anti. For differentiation to neutrophil-like cells (dHL-60), HL60 cells were cultured, for 5 days before any mRNA transfection, in IMDM with 10% FBS, 1% Anti-Anti and 1.25% DMSO, as described elsewhere with minor modifications^8^. These cells were cultured for 5 days with a medium change on the third day and have been designated as dHL60 cells.

### *In-vitro* transcription of mRNA

For *in-vitro* transfection of nano-luciferase mRNA, MEGAscript™ T7 Transcription Kit (Invitrogen, Thermo Fisher Scientific) was used, as per manufacturer’s instructions. The DNA of Nano-luciferase (both secretory and non-secretory expressing forms) was amplified from the plasmid pHIV-1 NL4·3Δenv-Luc and was inserted, downstream to the T7 promoter, between the 5’ untranslated region and 3’ untranslated regions followed by a poly-A tail and a T7 terminator in a pUC57 vector. The nano-luciferase plasmid was linearized using Kpn1 enzyme and purified using Gene Jet PCR purification kit (Thermo Fisher Scientific, USA). For *in-vitro* transcription, 1µg of linear DNA was used as a template. The reaction was set up with final concentrations of 7.5mM ATP, 7.5mM CTP, 7.5mM UTP, 1.5mM GTP, 6 mM Cap Analog, 1X Reaction Buffer and T7 Enzyme mix. Base-modified mRNA was prepared using one of the following bases, *N*^6^-methyladenosine (m6A), 5-methylcytidine (m5C), pseudo uridine (Ψ), and *N*^1^ methyl Ψ (Sapala Organics Private Limited, Hyderabad, India), which replaced their respective unmodified nucleosides at the same concentration. The GTP was added at lower concentration compared to other nucleotides due to presence of the Cap Analog in the ratio of 1:4 (GTP to Cap Analog), as per manufacturer’s instructions.

The DNA of eGFP was amplified and inserted, downstream to the T7 promoter, between the 5’ untranslated region and 3’ untranslated regions followed by a poly-A tail and a T7 terminator in a pETCON vector. The resulting plasmid was linearized using Kpn1 enzyme and purified using Gene Jet PCR purification kit (Thermo Fisher Scientific, USA). The *in-vitro* transcription was set up as before except for using a different Cap Analog (Sapala Organics Private Limited, Hyderabad, India) and 7.5 mM of GTP.

The reaction was set up at 37°C for 2.5 hours, after which DNase treatment was performed to remove any linear DNA contaminants. This was followed by lithium chloride precipitation to precipitate the RNA, which was then reconstituted in nuclease free water and stored at −80°C for transfections. In case of Cypridine luciferase mRNA, the modified mRNA was purchased from RNAVaxBio (Himachal Pradesh, India).

### mRNA transfection on mammalian and primary cells

#### Lipofectamine (endosomal) method

Cells were transfected using Lipofectamine 3000 (Invitrogen, Thermo Fisher Scientific), following the manufacturer’s instructions. RAW or dHL60 cells were transfected with 800ng of nano-luciferase mRNA (the non-secretory expressing form) in a 24-well plate with P3000 and reduced serum media, OptiMeM (Gibco, Thermo Fisher Scientific). The nano-luciferase transfections were done in duplicates and the average luminescence reading was tabulated for each time point. For Cypridine luciferase and Nano-luciferase (secretory expressing form) transfection, 1.5µg was transfected in 12-well plate with P3000, OptiMeM and luminescence was recorded once for each independent set. In the case of primary bone marrow derived macrophages (BMDM), transfection was done with a similar protocol using 2µg mRNA for cells in a 12-well plate. For BMDM transfection, the nano-luciferase mRNA transfections were done in a single set for each time point with cells isolated from three different mice. However, for Cypridine luciferase transfections, BMDMs were isolated from two mice and two sets of experiments were done from cells isolated from each mouse in 24-well plates with 800ng of mRNA.

#### Nucleofection (non-endosomal) method

Nucleofection is an advanced form of electroporation involving the delivery of nucleic acids to the nucleus and the cytoplasm of the recipient cells^9^. Nucleofection was done using Lonza Nucleofector^TM^ 2b with Cell Line Nucleofector kit V (Lonza Biosciences). Briefly, 1 million RAW 264.7 cells or dHL60 cells were resuspended in 82µl of Nucleofector solution and 18µl Supplement Solution to make a 100µl suspension. To this cell suspension, 3µg of unmodified and modified nano-luciferase mRNA was added in an aluminum nucleofector cuvette (Lonza). The cuvette was then nucleofected in the Nucleofector^TM^ 2b machine by using preset programs of D-032 (for RAW cells) and T-019 (for dHL60 cells). The cells were then recuperated in 1ml growth media and added to one well of a 24-well plate for each time-point.

For the primary bone marrow derived macrophages (BMDM) transfection, a similar protocol was followed using Mouse Macrophage Nucleofector Kit (Lonza). One million cells were resuspended in 82µl Nucleofector solution and 18µl Supplement solution, along with 4µg of mRNA in a Lonza nucleofection cuvette. The Lonza Nucleofector^TM^ 2b was set to the pre-set program Y-001 for BMDM. The cells were resuspended in 1ml media and divided into two wells of a 12-well plate for luciferase detection at two time points of 6h and 24h.

### Murine BMDM isolation, culture

All the animal experiments were performed in accordance with the Control and Supervision Rules, 1998 of the Ministry of Environment and Forest Act (Government of India), and the Institutional Animal Ethics Committee, Indian Institute of Science. Experiments were approved by the Committee for Purpose and Control and Supervision of Experiments on Animals (protocol number IISc CAF/ethics/718/2019). The animals were procured from Hylasco Bio-Technology Pvt. Ltd. (a Charles River Laboratories Subsidiary, Hyderabad, India). For BMDM isolation, C57BL6 mice were euthanized, and the bone marrow from the femur and tibia was isolated. The bone marrow was flushed to dislodge cells into 1X PBS and was passed through a mesh to make a uniform suspension. The cell suspension was centrifuged at 400x g for 4 minutes to pellet the cells. The pellet was resuspended in RBC lysis buffer, incubated for 5 minutes, and centrifuged at 400x g for 4 minutes to isolate cells free of RBC contaminants. For BMDM culture, the cells were cultured in DMEM with 10% FBS and 1.25μM β-mercaptoethanol. After two hours the media was replaced to remove the unbound cells, and the adherent cells were cultured in DMEM with 10% FBS and 1.25μM β-mercaptoethanol and 40ng/ml of GM-CSF (Granulocyte-macrophage colony stimulating factor-R&D Systems). On day two, the media was replaced with fresh media and 20ng/ml of GM-CSF. On day four, only half the media was replaced with fresh media and 20ng/ml GM-CSF. On day six, the cells were scraped and seeded in a 12-well plate for transfection on the following day, with growth media devoid of GM-CSF.

### Nano-Luciferase and Cypridine luciferase assay

For nano-luciferase assay, the cells were harvested and resuspended in 50µl 1X PBS and transferred to 96-well black plate (SPL Life Sciences, South Korea). For the assay, Nano-Glo Luciferase Assay System (Promega, USA) was used. To make the nano-luciferase assay reagent, one volume of assay substrate was mixed with 50 volumes of assay buffer. Then an equal volume of nano-luciferase assay reagent was added to the cell suspension, and luminescence was read in a microplate reader (Tecan) immediately. To detect the secreted form of the mRNA, equal volumes of cell supernatant and assay reagent were added for luminescence detection.

For Cypridine luciferase, Pierce Cypridina Luciferase Flash Assay Kit (Thermo Fisher Scientific) was used for luciferase detection. The supernatants of transfected cells were collected and 20µl was transferred to 96-well black plate (SPL Life Sciences). A luciferase working solution was made using Cypridina Flash Assay buffer and 1X Vargulin (final concentration), of which 50µl was added to the supernatants in each well. The luminescence was detected immediately in the microplate reader (Tecan).

### RNA isolation and RT-PCR

Nano-luciferase mRNA (unmodified and modified) transfected RAW 264.7 cells were subjected to RNA isolation to detect the mRNA levels of cytokines, TNF-α, IL-1β and IL-6. For this, the cells were transfected (by either Lipofectamine (endosomal) or Nucleofection (non-endosomal)) in a 12-well plate and incubated with the nano-luciferase mRNA for 3h, 6h and 24h. The cells were then lysed using Tri reagent (Sigma-Aldrich) and RNA was isolated using the RNeasy Mini kit (Qiagen) as per manufacturer’s instructions. After RNA isolation, the RNA was reverse transcribed to cDNA using iSCRIPT cDNA synthesis kit (Bio-Rad). The cDNA was then used to quantify the expression levels of the cytokines by using TB Green Premix Ex Taq II (Tli RNase H Plus) (Takara Bio Inc., Japan) TNF-α, IL-1β and IL-6 CFX96 Touch Real-Time PCR Detection System (Bio-Rad, USA). The sequences of the primers used, are provided in Supplementary Table S1. Comparative Ct method with normalization using 18s rRNA, was used for relative quantification of the genes.

### Enzyme Linked Immunosorbent Assay (ELISA)

To quantify the cytokine protein levels, ELISA was done from the cell supernatants of RAW and BMDM transfected cells. For this, the Mouse Uncoated TNF-α ELISA kit and Mouse Uncoated IL-6 ELISA kit were used (Invitrogen, Thermo Fisher Scientific). Initially, the capture antibodies were coated onto Maxi-binding 96-well transparent plates overnight at 4°C. The following day the plates were washed with 1X PBST thrice and blocked with ELISA Diluent (Invitrogen, Thermo Fisher Scientific) for one hour at room temperature. This was followed by incubation of 100µl cell supernatant in duplicates in the washed wells for two hours at room temperature. After two hours, the cell supernatants were discarded, and the plates were washed with 1X PBST thrice. Then the plates were coated with detection antibody for one hour at room temperature followed by subsequent washes and incubation with HRP-coupled enzyme for 30 minutes. The plates were washed rigorously and subjected to signal detection using TMB substrate. The reaction was stopped with 2N H_2_SO_4_ and the absorbance was detected at 450nm and 570nm using a microplate reader (Tecan).

### MTT assay

To determine viability of transfected RAW 264.7 cells, an MTT assay was done. For this, RAW246.7 cells (1X10^4 cells seeded in 48-well plate) were either transfected with N1 methyl Ψ modified and unmodified nano-luciferase mRNA using Lipofectamine 3000 as described earlier, or a similar number of cells were nucleofected with the mRNA and seeded in the 48-well plate. The transfected cells were incubated for 48 hours. Untransfected cells were kept as negative control while DMSO treated cells were kept as positive control. After 48 hours, the cells were washed with 1X PBS, and the cells were incubated with 20µl of MTT solution in 100µl DMEM for 4 hours at 37°C. The MTT solution was prepared in 1X PBS at a concentration of 5mg/ml. After 4 hours, formazan crystals formed on the 48-well plate were dissolved in 200µl DMSO, and the absorbance was measured using the microplate reader (Tecan) at 565nm. The experiment was done in duplicates and the viability was calculated as a percentage of cells alive as compared to untreated negative control.

### Flow cytometry for eGFP expression

To detect the expression of eGFP mRNA in transfected RAW cells, flow cytometry was done on BD FACSCelesta™ (Becton Dickinson, USA) and analyzed using De Novo FCS Express Software (Pasadena, CA, USA).

### Statistical analysis

At least three independent repeats were performed for all experiments, unless mentioned otherwise. One-way ANOVA with post-hoc Tukey test, or Kruskal-Wallis test with uncorrected Dunn’s test, or Welch’s ‘t’ test were performed to determine statistical differences in our data, and the specific tests have been mentioned in the respective figures. Statistically significant differences are represented as *p < 0.05, **p < 0.01, ***p < 0.001, and ****p < 0.0001. Data are presented as arithmetic mean ± standard deviation.

## Results

### Non-endosomal delivery results in higher mRNA expression in mammalian immune cell lines

To study the effect of mode of mRNA delivery and the need for modified mRNA, we measured the expression of unmodified and *N*^1^ methyl pseudo uridine (*N*^1^ methyl Ψ) modified mRNA, delivered through either the endosomal or non-endosomal routes in hard-to-transfect immune cell lines. In both the mouse macrophage and human neutrophil-like cell lines, mRNA expression was higher when the delivery was through the non-endosomal route as compared to the endosomal route (Figure 1A and 1B). The presence of *N*^1^ methyl Ψ modification of mRNA did not significantly affect expression if the delivery was non-endosomal, but for the endosomal delivery route, modified mRNA was required for high expression (Supplementary Table S2, S3). This observation is consistent with earlier studies which show better transfection efficiency rates with nucleofection (non-endosomal delivery)^10,11,12^. Importantly, the non-endosomal route of delivery did not result in the increased mRNA expression of inflammatory cytokines (TNFα, IL-1β and IL-6) in the mouse macrophage cell line when both unmodified and modified mRNA were used (Supplementary Figure S1). Additionally, neither method was detrimental for the survival of the cells (Supplementary Figure S2). We also observed a difference in expression levels between the two cell lines, which could be due to different immune cells having different capacities to take up and express foreign mRNA.

**Figure 1:**
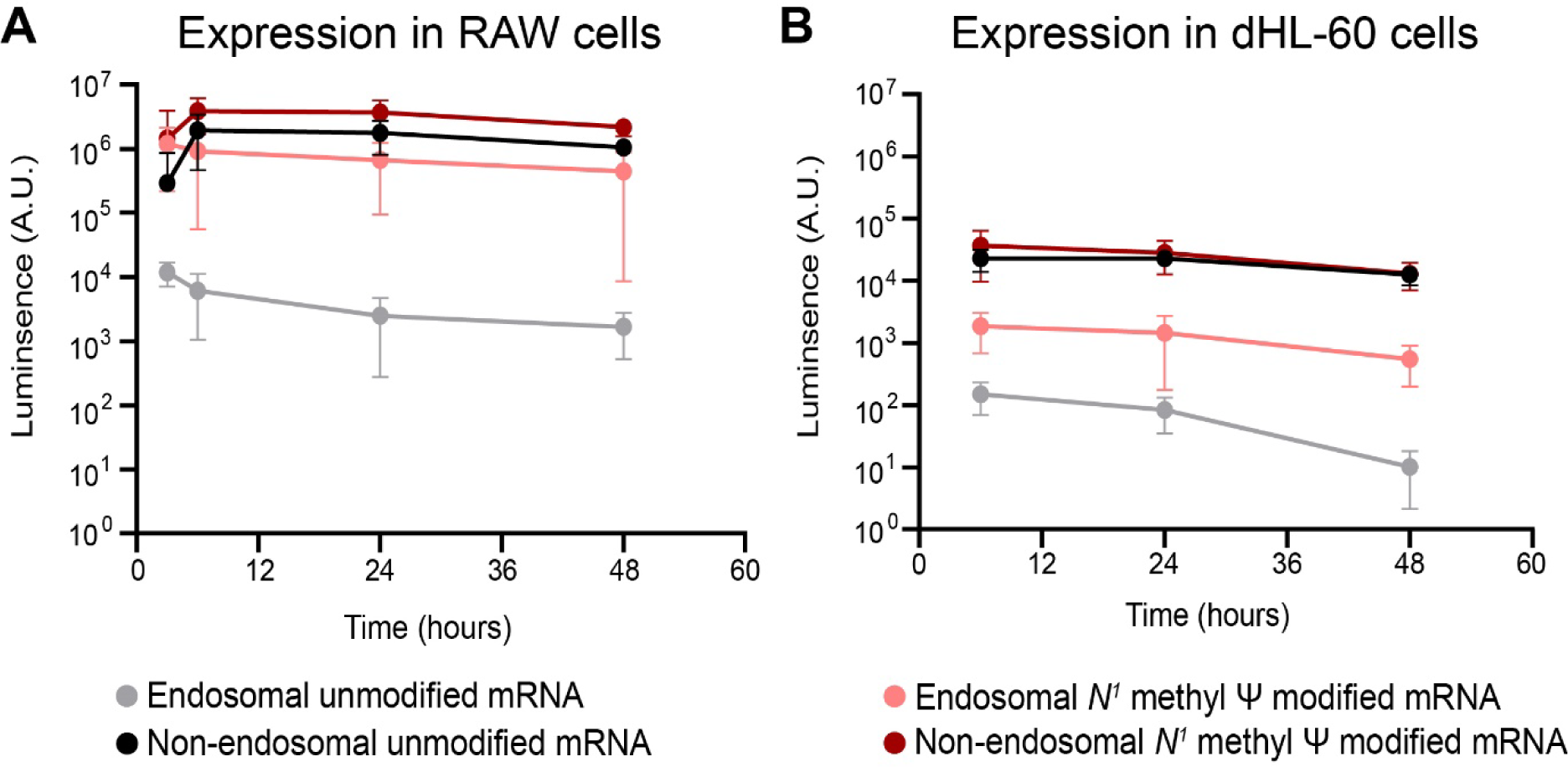
Effect of endosomal and non-endosomal delivery on mRNA expression in immune cell lines. **A:** Nano-luciferase expression in RAW 264.7 cells following transfection with unmodified or N1 methyl Ψ modified mRNA through either the endosomal or non-endosomal delivery route after 3, 6, 24 and 48 hours. (N=4). **B:** Nano-luciferase expression in dHL-60 cells following transfection with unmodified or N1 methyl Ψ modified mRNA through either the endosomal or non-endosomal delivery route after 6, 24 and 48 hours. (N= 3-4) Data are represented as mean ± S.D. Statistical test results are provided in supplementary table S2 and S3.

### Type of base-modification does not dictate mRNA expression and immune cell activation when route of delivery is non-endosomal

To confirm the hypothesis that modification was necessary only if the delivery was endosomal, we used four standard base modifications (*N*^6^-methyladenosine (m6A), 5-methylcytidine (m5C), pseudo uridine (Ψ), *N*^1^ methyl Ψ) of the mRNA and tested the expression levels when these were delivered through endosomal or non-endosomal routes. In the mouse macrophage cell line, we observe that when delivery was non-endosomal, the expression of unmodified mRNA was comparable to that of all modified versions of mRNA (except m6A, which resulted in lower expression), at two different times (Figure 2A and B). When the delivery was endosomal, Ψ and *N*^1^ methyl Ψ did indeed significantly increase expression (Supplementary Table S4), but the modifications did not play a role when the delivery was non-endosomal (Supplementary Table S5). These data confirm the hypothesis that base-modifications are not necessary when delivering through the non-endosomal route, but N1 methyl Ψ modification is important to obtain high expression when delivering through the endosomal route.

**Figure 2:**
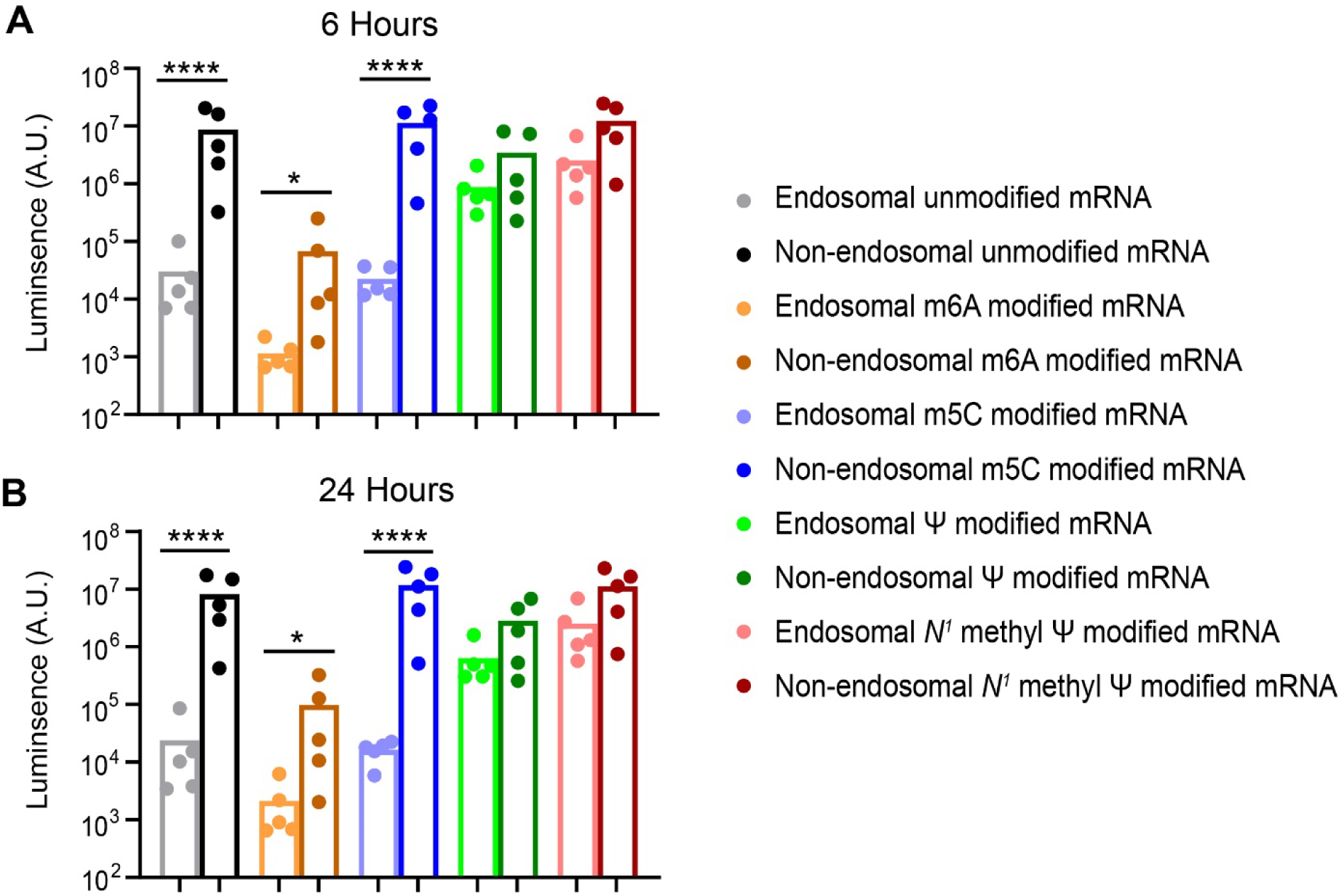
Influence of delivery route and mRNA modifications on expression in RAW (murine macrophage) cell line. Nano-luciferase expression in RAW 264.7 cells following transfection with unmodified and four modified versions (*N*^6^-methyladenosine (m6A), 5-methylcytidine (m5C), pseudo uridine (Ψ), *N*^1^ methyl Ψ) of mRNA through either the endosomal and non-endosomal delivery route after 6 hours **(A)** and 24 hours **(B)**. (N=5) Statistical testing was performed on the logarithm values of luminescence using one-way ANOVA followed by Tukey post-hoc test. * p < 0.05, *** p < 0.0001 and not significant where nothing is indicated. Additional statistical comparisons are available in Supplementary table S4 and S5.

Next, to determine how the mode of mRNA delivery and the base-modifications affect the activation of cells, we measured the secretion of two inflammatory cytokines TNF-α and IL-6. Unlike mRNA expression levels, IL-1β protein levels were not detectable and hence the data is not shown. The non-endosomal delivery route resulted in lower TNF-α secretion when compared to the endosomal delivery route when unmodified, m6A modified and *N*^1^ methyl pseudo uridine modified versions of the mRNA were used at both 6 and 24 hours post transfection (Figure 3A).

**Figure 3:**
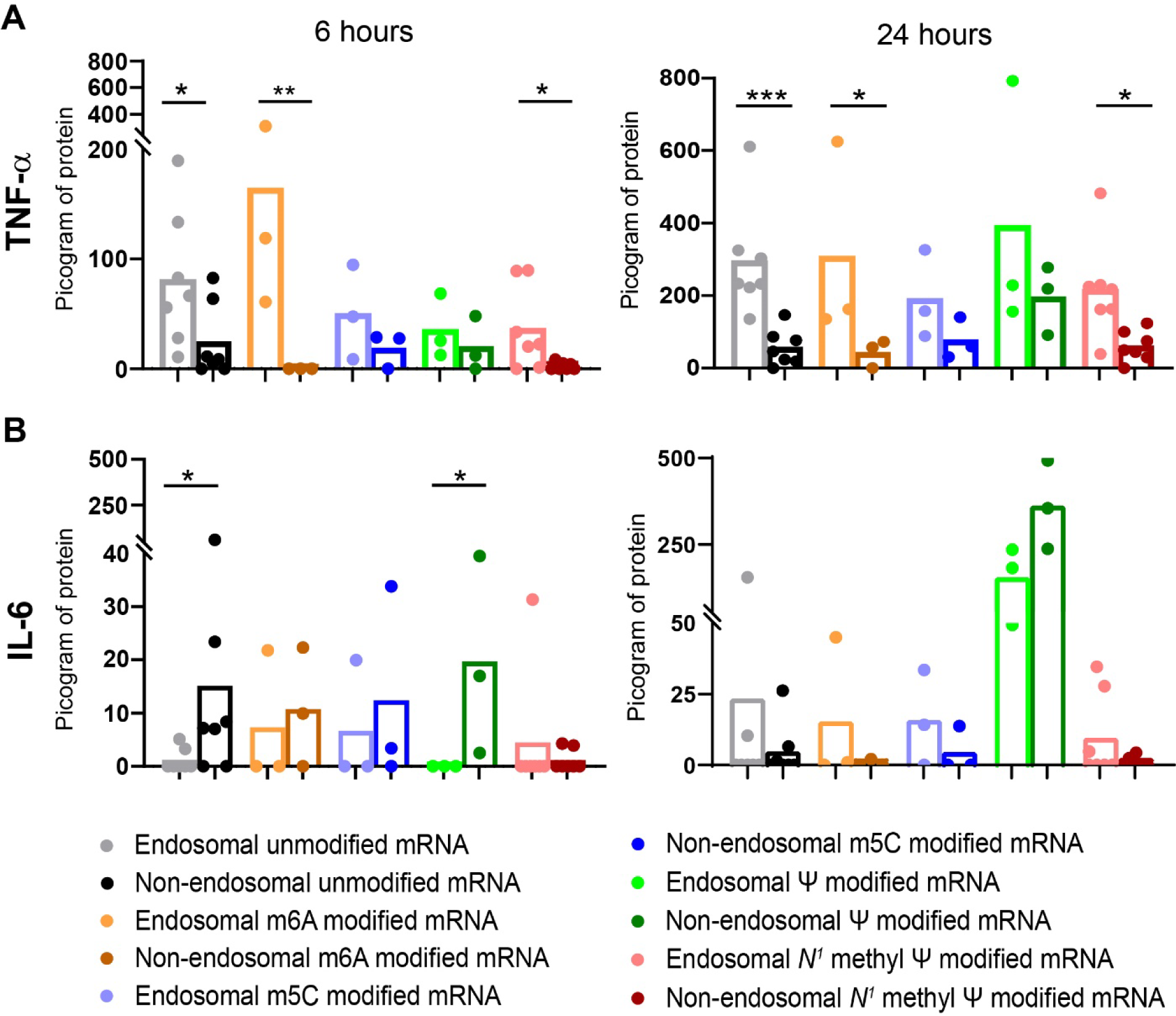
Absence of inflammatory cytokine secretion upon non-endosomal delivery of mRNA. (**A**) TNF-α protein and (**B**) IL-6 protein expression levels (ELISA) in RAW 264.7 cell supernatants that were transfected with unmodified or modified mRNA through either the endosomal or non-endosomal delivery route after 6 hours (*left panel*) and 24 hours (*right panel*). (N=3-7) Statistical testing was performed using the Kruskal Wallis Test. * p < 0.05, ** p < 0.01, *** p < 0.0001 0001 and not significant where nothing is indicated. Additional statistical comparisons are available in Supplementary table S6 and S7.

When comparisons were made among only the endosomal delivery route (different modifications) or among only non-endosomal delivery route (different modifications), no significant difference was observed (Supplementary Table S6 and S7). Surprisingly, at 6 hours, IL-6 secretion was increased in the non-endosomal delivery route as compared to the endosomal, upon the use of unmodified and pseudo uridine modified mRNA (Figure 3B). However, N1-methyl pseudo uridine modified mRNA resulted in no significantly detectable secretion of IL-6 at both 6 and 24 hours post-transfection. At 24 hours post non-endosomal delivery, the IL-6 levels in the unmodified mRNA group also dropped and were not different than other modifications apart from the pseudouridine modification (Supplementary Table S8 and S9). These data show that a differential response in TNF-α and IL-6 secretion is observed when using unmodified mRNA and delivering it through the non-endosomal route as compared to the endosomal route. However, the use of *N*^1^-methyl pseudo uridine modification ensures that secretion of both these inflammatory cytokines remains undetectable.

### Primary murine macrophage cells also show higher expression and lower cellular activation upon non-endosomal delivery

To validate the effect of non-endosomal delivery on expression and cellular activation, primary mouse macrophages were used. These primary cells were isolated from the mouse bone marrow and mRNA expression was studied following endosomal or non-endosomal delivery using unmodified and *N*^1^ methyl pseudo uridine modified mRNA. Consistent with the cell line data, unmodified mRNA shows higher expression when delivered through the non-endosomal compared to the endosomal route, in primary macrophages (Figure 4A). The *N*^1^-methyl Ψ modified mRNA did not have any significant differences in expression when either mode of delivery was used. Importantly, the non-endosomal delivery route resulted in no secretion of the inflammatory cytokines, TNF-α and IL-6, when either unmodified or modified mRNA was used, however, the endosomal delivery route required modifications to not activate the cells (Figure 4B and 4C).

**Figure 4:**
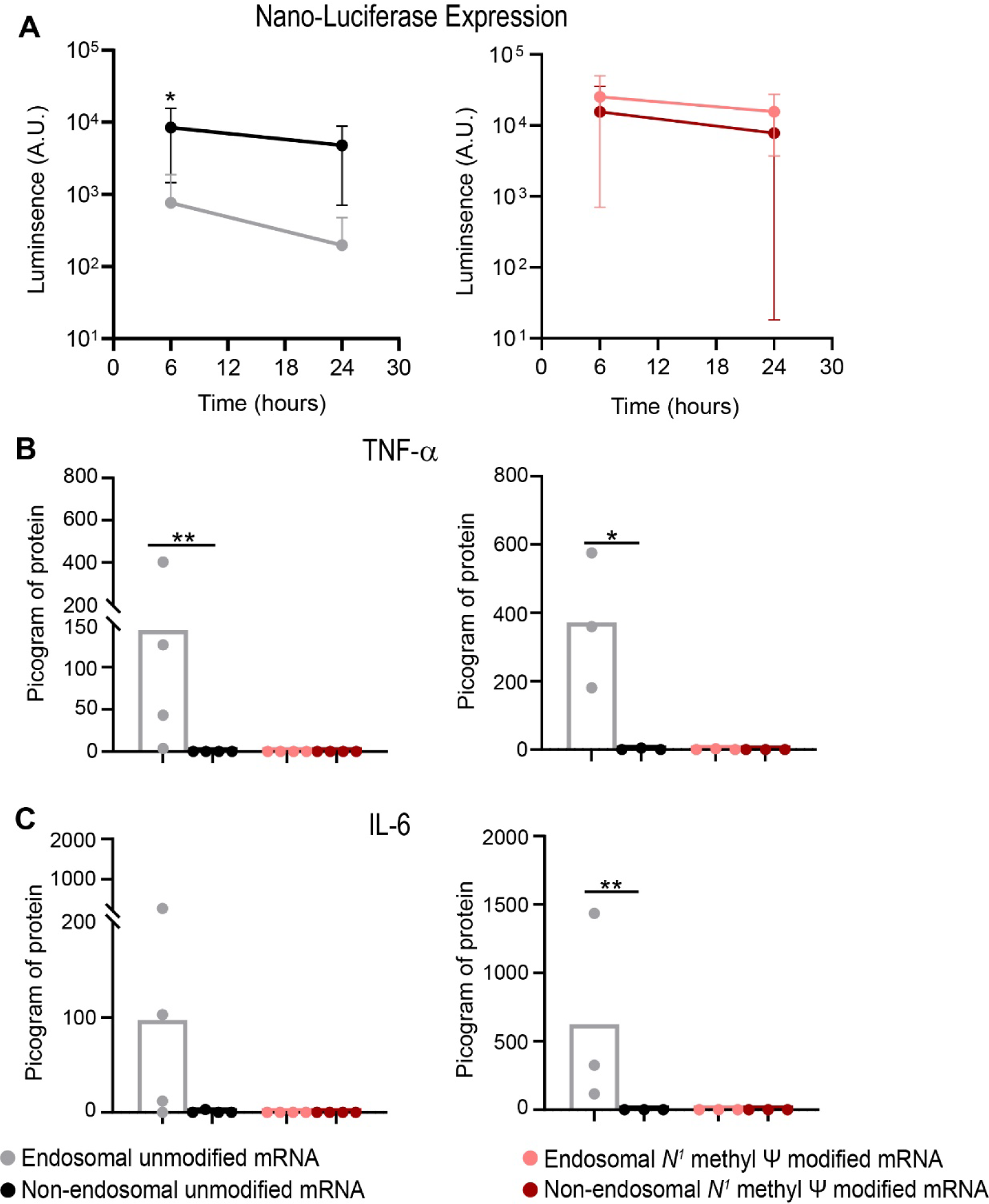
mRNA expression and inflammatory response in mouse bone marrow derived macrophages (BMDM). **A:** Nano-luciferase expression in BMDM cells following transfection with unmodified (*left*) or *N*^1^ methyl Ψ modified (*right*) mRNA through either the endosomal or non-endosomal delivery route after 6 hours (N=4) and 24 hours (N=3). Welch’s ‘t’ test was used and * p <0.05. (**B**) TNF-α protein and (**C**) IL-6 expression levels (ELISA) in BMDM cell supernatants that were transfected with unmodified or modified mRNA through either the endosomal or non-endosomal delivery route after 6 hours (*left panel, N=4*) and 24 hours (*right panel, N=3*). Kruskal Wallis multiple comparison test was used and * p < 0.05, ** p < 0.01.

### Sustained expression following non-endosomal delivery

Next, we addressed the question of whether delivery by endosomal and non-endosomal routes would result in sustained expression of mRNA. We used a secreted version of the nano-luciferase mRNA to study sustenance of expression in RAW macrophages, and observed that delivery of modified mRNA *via* the endosomal route and both unmodified and modified mRNA *via* the non-endosomal route resulted in higher expression until 72 hours post-delivery when compared to the unmodified mRNA delivered *via* the endosomal route (Figure 5). These results are consistent with previous studies on sustained expression following delivery of modified mRNA^3,5^.

**Figure 5:**
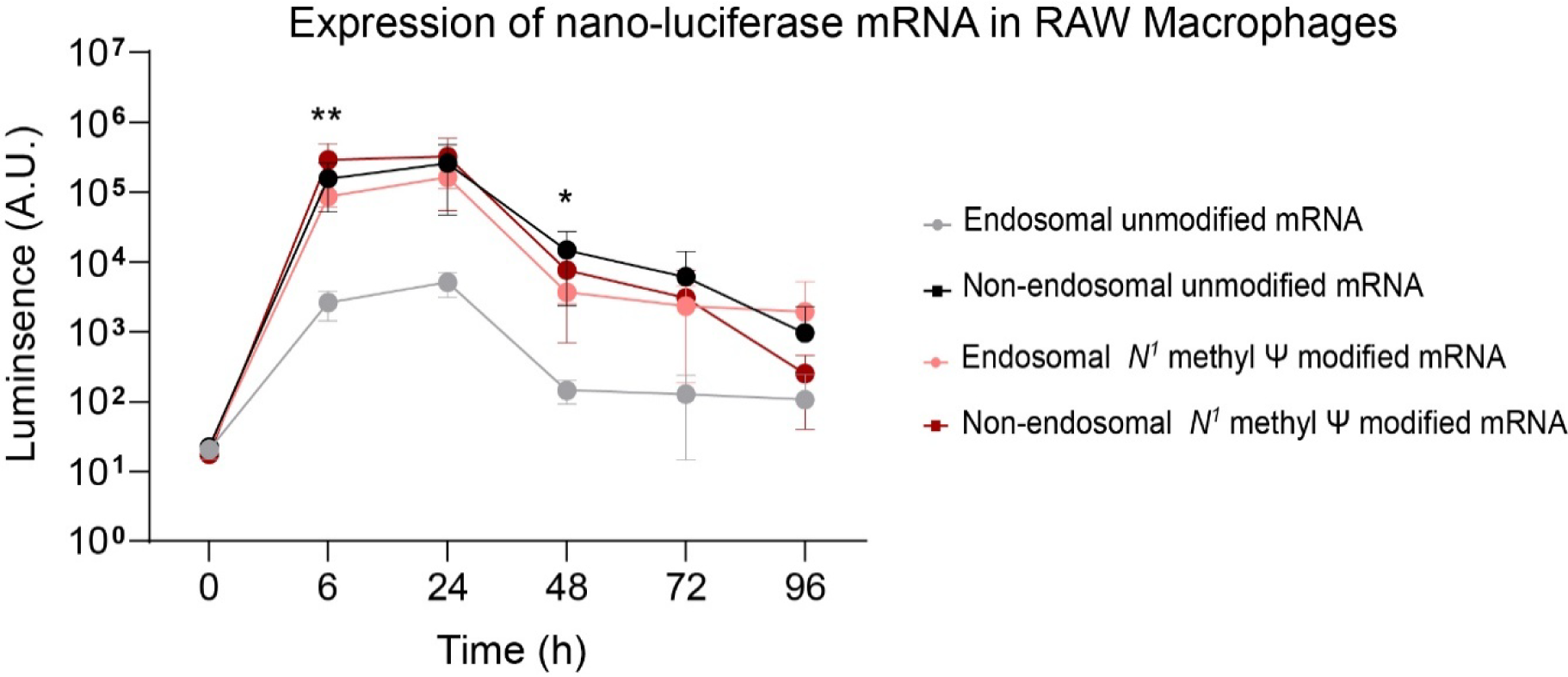
Sustained expression. Nano-luciferase expression in RAW 264.7 cells following delivery of unmodified or *N*^1^-methyl Ψ modified mRNA either through the endosomal or non-endosomal delivery route after 6, 24, 48, 72 and 96 hours. Data are based on N=3 independent experiments and are represented as mean ± S.D. P-values were calculated using a One Way ANOVA (log-normalized data) at each time point, and * p<0.05 and ** p<0.01.

### Delivery of different mRNA

One question that remains unaddressed is if the aforementioned observations are specific to a type of mRNA used in our experiments. To confirm that the observed effects were not due to a specific type of mRNA used for all previous studies (nano-luciferase mRNA), we utilized an mRNA that codes for enhanced green fluorescence protein (eGFP). The advantage of using this specific mRNA is that following transfection, flow cytometry may be performed to determine expression of mRNA at a single cell level. Unmodified or *N*^1^-methyl Ψ modified mRNA was transfected in RAW macrophages using the endosomal or non-endosomal routes, and protein expression was monitored using flow cytometry at 6 hours and 24 hours post transfection. This specific mRNA showed poor expression when delivered through the endosomal route. However, protein expression was observed when the non-endosomal delivery route was used, and importantly, the expression levels of both unmodified and modified mRNA were similar when using this route of delivery (Figure 6 and Supplementary Fig S3).

**Figure 6:**
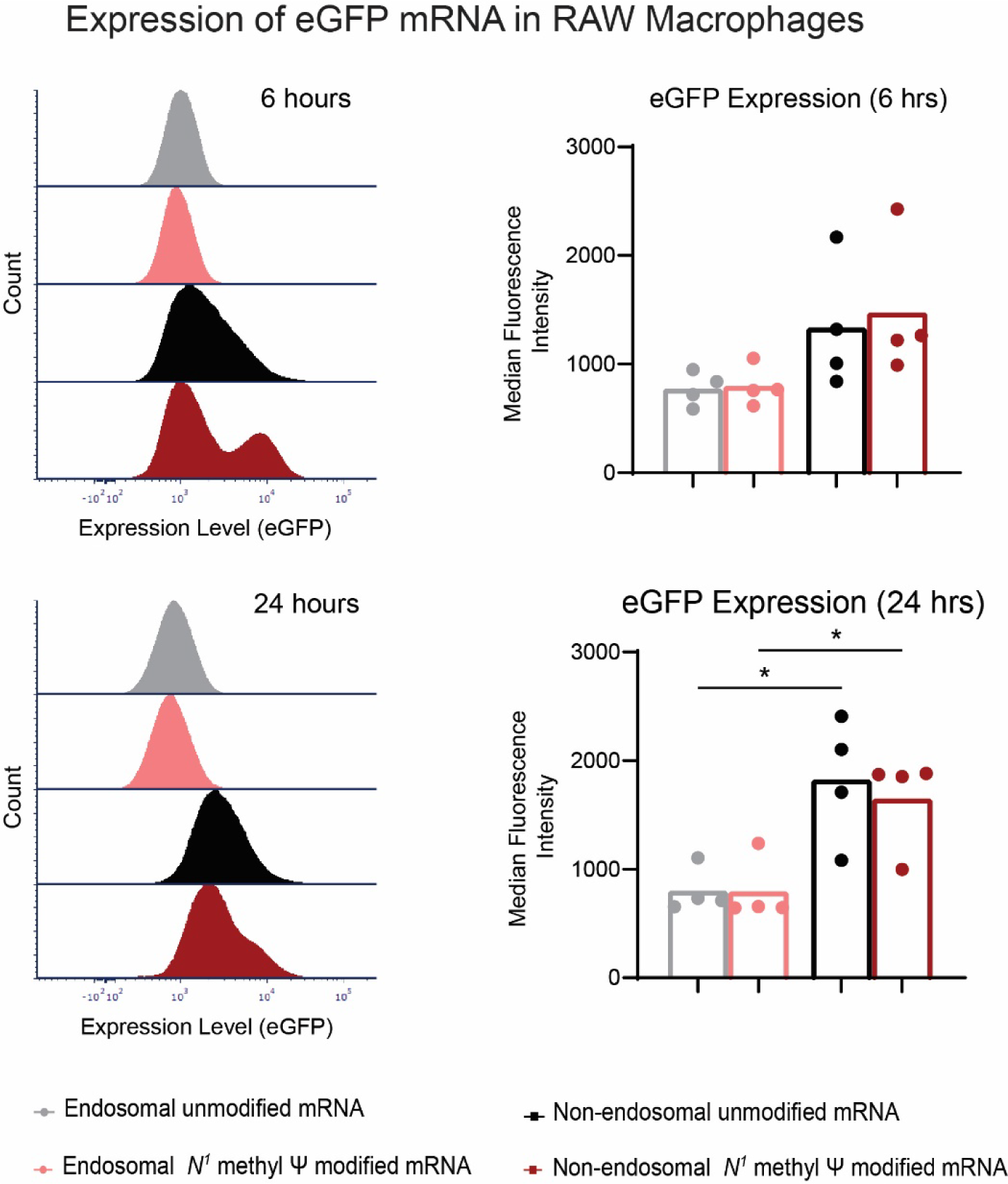
Expression of a different reporter mRNA (enhanced green fluorescence protein – eGFP). Expression of unmodified and *N*^1^-methyl Ψ modified eGFP mRNA in RAW 264.7 cells that were delivered through the endosomal or non-endosomal routers after 6 (top panel) and 24 (bottom panel) hours as detected by flow cytometry. In the left are histogram plots depicting expression of the eGFP, and on the right is a plot of the median fluorescence intensity values. Data are from N=4 independent experiments and are represented as mean ± S.D. P-values were calculated by utilizing One Way ANOVA. Data is non-significant at 6 hours, and * p<0.05 for specific comparisons at 24 hours.

As all the previous studies were performed using mRNA that was prepared in our laboratory, we also verified our results of mRNA expression in immune cells when delivery is *via* the non-endosomal route, by using a commercially available mRNA (mRNA coding for Cypridine luciferase). In the studies using the purchased mRNA, we observe that the non-endosomal delivery route can sustain expression of mRNA for the same time duration as endosomal delivery. Additionally, using the *N*^1^-methyl Ψ modified version of this mRNA, we observed that non-endosomal delivery results in enhanced expression as compared to the endosomal delivery in both the mouse macrophage cell line and primary mouse bone-marrow derived macrophages (Supplementary Figure S4).

## Discussion

With the success of COVID mRNA vaccines, there is a surge in interest on mRNA-based therapeutics. In such therapeutics, it is assumed that nucleoside modification, such as the *N*^1^-methyl Ψ modification, is necessary. However, *N*^1^-methyl Ψ modified mRNA is not very efficient when delivered directly *in vivo*. To further protect the mRNA from degradation, facilitate their uptake by specific cells in the body, and possibly enable sustained release, thereby enabling efficient *in vivo* expression of mRNA, lipid nanoparticles (LNPs) have been described to be essential^13,14,15^.

However, both base-modification and the use of LNPs add to the cost and complexity of developing mRNA-based therapeutics. For instance, LNPs utilize the endosomal pathway for delivery and consequently “endosomal escape” of mRNA at a pharmacologically relevant level, becomes a matter of further optimization^16,17,18^. While the need for both base-modification and LNPs when the mRNA-therapeutic needs to be delivered *in vivo* is unquestioned, it remains unclear if one or both are required for *ex vivo* cell engineering. When mRNA is being delivered to cells *ex vivo*, in culture, LNPs can be avoided through the use of electroporation or nucleofection. Nucleofection can deliver mRNA to different cell types with improved expression and lower toxicity^19^. It has been observed that nucleofection helps to deliver nucleic acids to primary human T cells^20^, human pre-B cells^21^ as well as macrophages^22^ efficiently, suggesting that non-endosomal delivery can facilitate delivery of mRNA to a wide variety of cell types, especially immune cells, which in general are difficult to transfect. The question that remains is when using non-endosomal delivery for *ex vivo* delivery of mRNA, is base modification necessary.

To address this question, here we compared the efficacy of expression and immune reactivity of unmodified and base-modified mRNA, delivered either through the endosomal or non-endosomal route. Our data shows that non-endosomal delivery of mRNA can lead to efficient protein expression, irrespective of nucleoside modifications. In the cell lines and primary immune cells that we evaluated, base modification was not necessary for high and sustained protein expression, when using the non-endosomal delivery route. Also, consistent with previous reports^5,23,24,25^, we demonstrated that *N*^1^-methyl Ψ mRNA modification was necessary for higher expression and lower inflammatory responses in the same immune cells if an endosomal delivery route was used.

In their landmark paper in 2005, Kariko et al. demonstrated that base-modified mRNA (that are present in mammalian cells but not in bacteria) have a lowered activation of innate immune responses^1^. This paper laid the foundation for the use of base-modified mRNA, and using such modified mRNA resulted in better *in vitro* and *in vivo* expression of the coded protein. However, this and many other subsequent papers, including Andries et al. who showed that *N*^1^-methyl Ψ modification was superior for expression and had lower immunogenicity^5^, have used a variant of liposome formulation or lipid nanoparticles for the delivery of the modified mRNA. Our results are consistent with all these studies, and we show that if mRNA is delivered using lipofectamine (endosomal delivery route), then base modification is necessary for higher expression and lower immunogenicity.

However, we demonstrate that such base modifications may be unnecessary if the mRNA is delivered through a non-endosomal route (electroporation) that potentially results in direct cytoplasmic delivery. Direct cytoplasmic delivery may be useful for *ex vivo* cell engineering^26^. Our data on equally high expression of unmodified mRNA (as compared to modified mRNA) that is delivered through the non-endosomal route suggests the following. First, previous work demonstrating that specific RNA modifications that promote structural stability result in superior translation efficiency^27^, may have promoted stability of the mRNA in the formulation and might not be necessary if direct cytoplasmic delivery is performed. Second, for *ex vivo* cell engineering, the requirement of new ionizable lipids^28,29^ or molecules that promote endosomal escape^30^ are likely to be unnecessary.

A noteworthy point is that a transient increase in IL-6 secretion by RAW macrophages was observed at early time points post transfection (figure 3B) when using the unmodified mRNA, but the same was not observed in primary mouse macrophages (figure 4C), suggesting that unmodified (and *N*^1^-methyl Ψ modifed) mRNA is likely to be non-immunogenic if delivered through a non-endosomal route. Interestingly, the pseudouridine modification (and not other modifications or unmodified version of the mRNA) resulted in an increased IL-6 expression regardless of the delivery route. Others have reported IL-6 secretion when pseudouridine modified mRNA is delivered through the endosomal route^7,31^. When mRNA is delivered directly into the cytoplasm, it is possible that the pseudouridine modification results in higher IL-6 secretion due to an altered interaction with the cytoplasmic pattern recognition receptors like the Retinoic acid-inducible gene (RIG)-I-like receptors (RLRs)^32^, but this remains to be verified.

## Conclusions

Our data provides further evidence in support of the claim that when delivering mRNA through an endosomal route, *N*^1^-methyl Ψ mRNA modification was better than several other nucleoside modifications in terms of mRNA expression and reducing the severity of inflammatory responses. We also demonstrate that both unmodified and base-modified mRNA result in equally high expression and low to absent inflammatory responses if a non-endosomal delivery route is used for *ex vivo* delivery of mRNA to specific immune cells. Together, our data suggests that non-endosomal delivery of mRNA might be an effective tool for *ex vivo* immune cell engineering, and in such an instance, base-modification will not be necessary.

## Supplementary Information

Supplementary Figure S1: transcript levels of cytokines in RAW cells

Supplementary Figure S2: assay on cell survival following delivery of mRNA

Supplementary Figure S3: flow cytometry gating strategy

Supplementary Figure S4: sustained expression of cypridine luciferase

Supplementary Table S1: list of primers

Supplementary Table S2-S9: detailed statistical analysis

## Acknowledgements

The work presented here was funded by the Ignite Life Science Foundation, India to RV and SJ. The authors also thank the Indian Institute of Science for providing bridge funding for some parts of this work. We acknowledge Shruthi Ksheera Sagar for assistance with nucleofection experiments. B.G. acknowledges the support provided by the national postdoctoral fellowship scheme of the Science and Engineering Board, Department of Science and Technology, Govt. of India (PDF/2022/001783). DC acknowledges Council for Scientific and Industrial Research, Govt. of India (09/079 (2837) / 2019-EMR-1) and MN acknowledges Ministry of Education, Govt. of India (PMRF ID:0201020) doctoral fellowships. RV is a JC Bose fellow of Department of Science and Technology, Govt. of India. SJ is an Intermediate fellow of the DBT-Wellcome India Alliance program.

## Notes

### Competing Interest Statement

The authors have declared no competing interest.

