## Supplementary for "*Ex Vivo* Delivery of mRNA to Immune Cells *Via* a Non-Endosomal Route Obviates the Need for Nucleoside Modification"

**Supplementary Figures S1 – S4:** **Page 2-5**

**Supplementary Table S1 – S9** **Page 6-15**

**Supplementary Figure S1:**

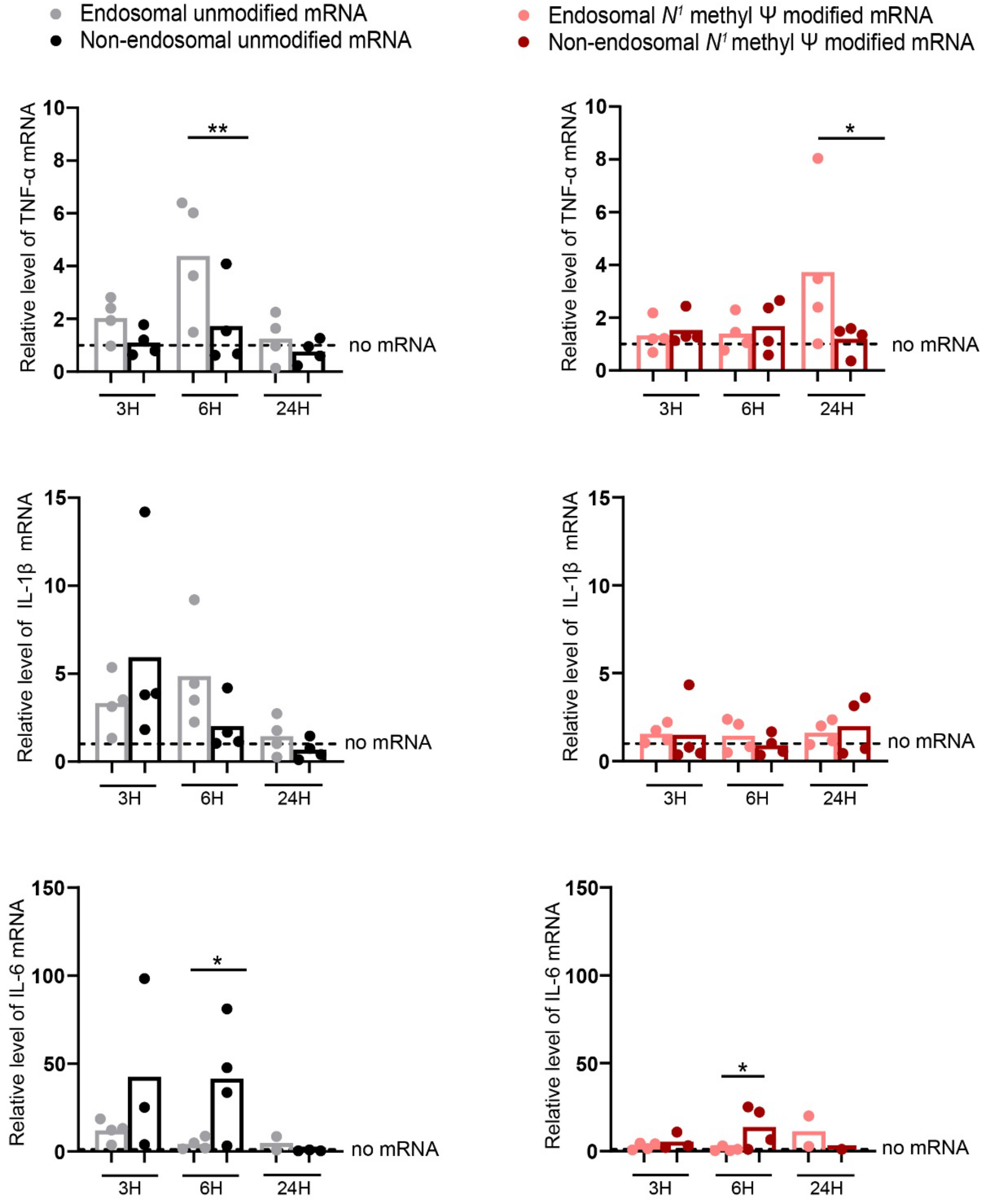


**Figure S1: mRNA levels of inflammatory cytokines in RAW264.7 cells transfected with unmodified or $N^1\Psi$ modified mRNA through endosomal or non-endosomal route.**

Quantitative PCR to show TNF-α, IL-1β and IL-6 mRNA levels in RAW cells after endosomal and non-endosomal delivery of unmodified (left panel) and $N^1$ methyl pseudo uridine ($N^1\Psi$) modified (right panel) mRNA, at 3, 6 and 24 hours post transfection. 18srRNA was used as a normalizing control. (N=4)

P-values were calculated by utilizing One Way ANOVA. ns: non-significant, *P < 0.05, **P < 0.01, ***P < 0.0001.

**Supplementary Figure S2:**

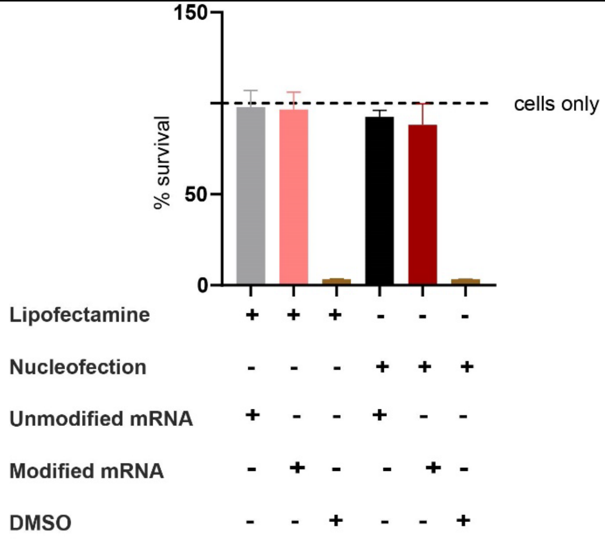


**Figure S2: Effect of mRNA delivery route and modification on viability of RAW macrophage cells.**

MTT assay to determine cell viability at 48 hours post transfection with unmodified and $N^1\Psi$ modified nano-luciferase mRNA delivered through the endosomal or non-endosomal route. N=3

Data are represented as mean ± S.E.M. DMSO was used as a positive control for cell killing.

**Supplementary Figure S3:**

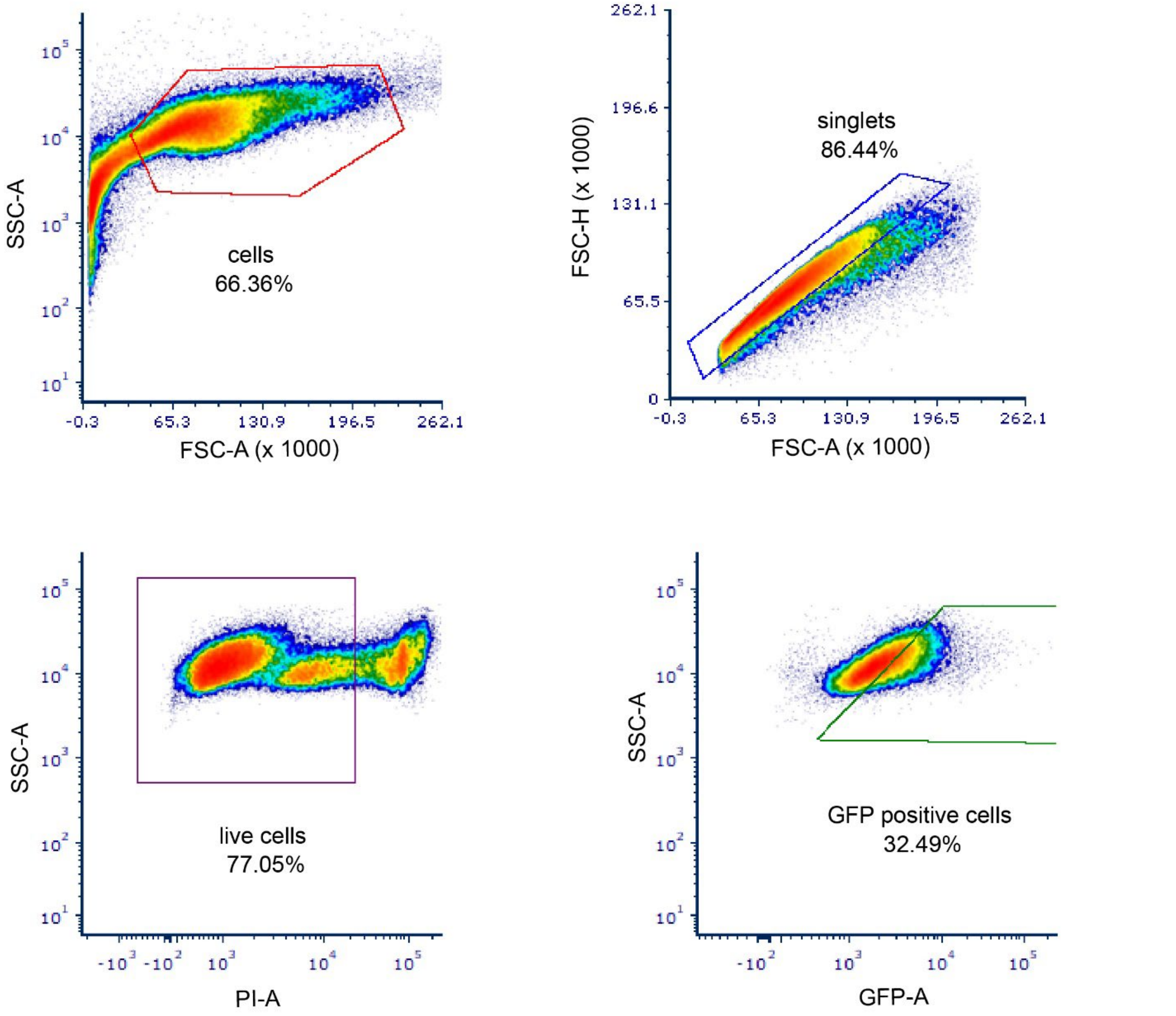


**Figure S3: Gating strategy utilized to determine expression of enhanced green fluorescence protein (eGFP) using flow cytometry.**

**Supplementary Figure S4:**

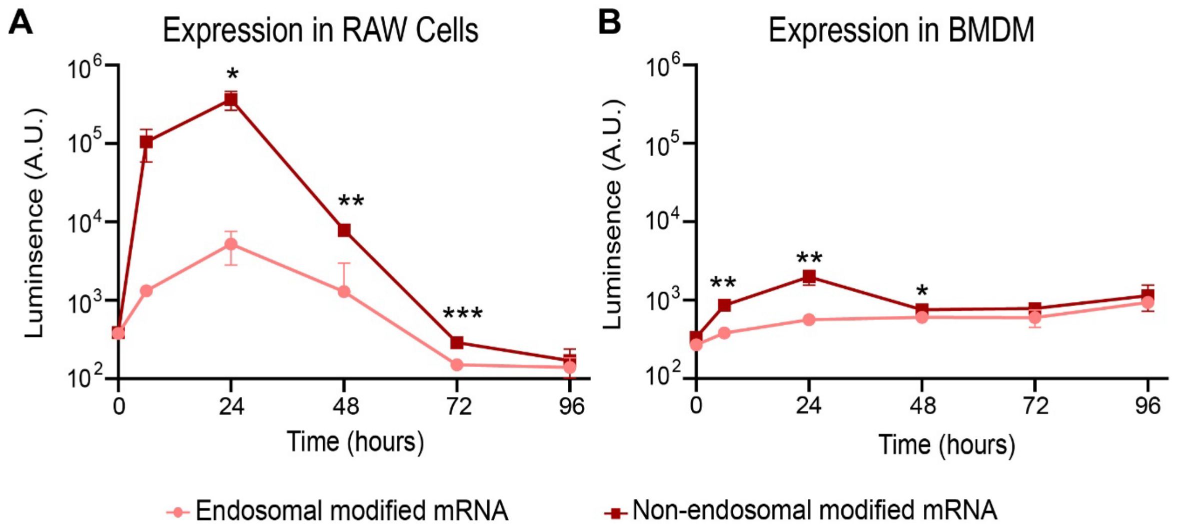


**Figure S4: Impact of delivery route on sustainability of mRNA expression.**
Cypridine luciferase expression in RAW cells (**A**) and BMDM cells (**B**) at different time points of 6, 24, 48, 72 and 96 hours. For RAW cells, N=3, for BMDM, N=4 (experiment done in duplicates from two mice bone marrow).

Data are represented as mean ± S.D. P-values were calculated by utilizing Two Way ANOVA. ns: non-significant, * $p<0.05$, ** $p<0.01$, *** $p<0.0001$.

**Supplementary Table S1: List of primers**

| Gene | Forward Primer | Reverse Primer |
|---|---|---|
| IL-1β | 5’-CAACCAACAAGTGATATTCTCCATG-3’ | 5’-GATCCACACTCTCCAGCTGCA-3’ |
| IL- 6 | 5’-TACCACTTCACAAGTCGGAGGC-3’ | 5’-CTGCAAGTGCATCATCGTTGTTC-3’ |
| Tnf-α | 5’-GGTGCCTATGTCTCAGCCTCTT-3’ | 5’-GCCATAGAACTGATGAGAGGGAG-3’ |
| 18S rRNA | 5’-GCAATTATTCCCCATGAACG-3' | 5’-GGCCTCACTAAACCATCCAA-3' |

**Supplementary Table S2: One-way ANOVA followed by post-hoc Tukey test (of log values of nano luciferase expression) to determine statistical significance between mRNA modification and delivery route in RAW264.7 cells at different time points (supporting data presented in figure 1A).**

**A. 3 hours**

| Tukey test (post ANOVA) | Summary | Adjusted P Value |
|---|---|---|
| Endosomal unmodified mRNA vs. Non-Endosomal unmodified mRNA | ns | 0.9606 |
| Endosomal unmodified mRNA vs. Endosomal $N^1$ methyl Ψ modified mRNA | * | 0.0327 |
| Endosomal unmodified mRNA vs. Non-endosomal $N^1$ methyl Ψ modified mRNA | ns | 0.1654 |
| Endosomal $N^1$ methyl Ψ modified mRNA vs. Non-Endosomal unmodified mRNA | ns | 0.0756 |
| Non-Endosomal unmodified mRNA vs. Non-endosomal $N^1$ methyl Ψ modified mRNA | ns | 0.3366 |

| | | |
|---|---|---|
| Endosomal $N^1$ methyl Ψ modified mRNA vs. Non-endosomal $N^1$ methyl Ψ modified mRNA | ns | 0.7711 |

B. **6 hours**

| Tukey test (post ANOVA) | Summary | Adjusted P Value |
|---|---|---|
| Endosomal unmodified mRNA vs. Non-Endosomal unmodified mRNA | **** | <0.0001 |
| Endosomal unmodified mRNA vs. Endosomal $N^1$ methyl Ψ modified mRNA | *** | 0.0001 |
| Endosomal unmodified mRNA vs. Non-endosomal $N^1$ methyl Ψ modified mRNA | **** | <0.0001 |
| Endosomal $N^1$ methyl Ψ modified mRNA vs. Non-Endosomal unmodified mRNA | ns | 0.6609 |
| Non-Endosomal unmodified mRNA vs. Non-endosomal $N^1$ methyl Ψ modified mRNA | ns | 0.6265 |
| Endosomal $N^1$ methyl Ψ modified mRNA vs. Non-endosomal $N^1$ methyl Ψ modified mRNA | ns | 0.1339 |

C. **24 hours**

| Tukey test (post ANOVA) | Summary | Adjusted P Value |
|---|---|---|
| Endosomal unmodified mRNA vs. Non-Endosomal unmodified mRNA | **** | <0.0001 |

| | | |
|---|---|---|
| Endosomal unmodified mRNA vs. Endosomal $N^1$ methyl Ψ modified mRNA | **** | <0.0001 |
| Endosomal unmodified mRNA vs. Non-endosomal $N^1$ methyl Ψ modified mRNA | **** | <0.0001 |
| Endosomal $N^1$ methyl Ψ modified mRNA vs Non-Endosomal unmodified mRNA | ns | 0.3148 |
| Non-Endosomal unmodified mRNA vs. Non-endosomal $N^1$ methyl Ψ modified mRNA | ns | 0.8046 |
| Endosomal $N^1$ methyl Ψ modified mRNA vs. Non-endosomal $N^1$ methyl Ψ modified mRNA | ns | 0.0775 |

**D. 48 hours**

| Tukey test (post ANOVA) | Summary | Adjusted P Value |
|---|---|---|
| Endosomal unmodified mRNA vs. Non-Endosomal unmodified mRNA | **** | <0.0001 |
| Endosomal unmodified mRNA vs. Endosomal $N^1$ methyl Ψ modified mRNA | **** | <0.0001 |
| Endosomal unmodified mRNA vs. Non-endosomal $N^1$ methyl Ψ modified mRNA | **** | <0.0001 |
| Non-Endosomal unmodified mRNA vs. Endosomal $N^1$ methyl Ψ modified mRNA | ns | 0.0847 |
| Non-Endosomal unmodified mRNA vs. Non-endosomal $N^1$ methyl Ψ modified mRNA | ns | 0.4616 |
| Endosomal $N^1$ methyl Ψ modified mRNA vs. Non-endosomal $N^1$ methyl Ψ modified mRNA | ** | 0.0062 |

**Supplementary Table S3: One-way ANOVA followed by post-hoc Tukey test (of log values of nano luciferase expression) to determine statistical significance between mRNA modification and delivery route in dHL-60 cells at different time points (supporting data presented in figure 1B).**

**A. 6 hours**

| Tukey test (post ANOVA) | Summary | Adjusted P Value |
|---|---|---|
| Endosomal unmodified mRNA vs. Non-Endosomal unmodified mRNA | *** | 0.0007 |
| Endosomal unmodified mRNA vs. Endosomal $N^1$ methyl Ψ modified mRNA | * | 0.0133 |
| Endosomal unmodified mRNA vs. Non-endosomal $N^1$ methyl Ψ modified mRNA | **** | <0.0001 |
| Endosomal $N^1$ methyl Ψ modified mRNA vs. Non-Endosomal unmodified mRNA | ** | 0.0044 |
| Non-Endosomal unmodified mRNA vs. Non-endosomal $N^1$ methyl Ψ modified mRNA | ns | 0.9489 |
| Endosomal $N^1$ methyl Ψ modified mRNA vs. Non-endosomal $N^1$ methyl Ψ modified mRNA | ** | 0.0021 |

**B. 24 hours**

| Tukey test (post ANOVA) | Summary | Adjusted P Value |
|---|---|---|
| Endosomal unmodified mRNA vs. Non-Endosomal unmodified mRNA | **** | <0.0001 |
| Endosomal unmodified mRNA vs. Endosomal $N^1$ methyl Ψ modified mRNA | ** | 0.0095 |

| | | |
|---|---|---|
| Endosomal unmodified mRNA vs. Non-endosomal $N^1$ methyl Ψ modified mRNA | **** | <0.0001 |
| Endosomal $N^1$ methyl Ψ modified mRNA vs. Non-Endosomal unmodified mRNA | ** | 0.0013 |
| Non-Endosomal unmodified mRNA vs. Non-endosomal $N^1$ methyl Ψ modified mRNA | ns | 0.9489 |
| Endosomal $N^1$ methyl Ψ modified mRNA vs. Non-endosomal $N^1$ methyl Ψ modified mRNA | ** | 0.0010 |

**C. 48 hours**

| Tukey test (post ANOVA) | Summary | Adjusted P Value |
|---|---|---|
| Endosomal unmodified mRNA vs. Non-Endosomal unmodified mRNA | **** | <0.0001 |
| Endosomal unmodified mRNA vs. Endosomal $N^1$ methyl Ψ modified mRNA | ** | 0.0095 |
| Endosomal unmodified mRNA vs. Non-endosomal $N^1$ methyl Ψ modified mRNA | **** | <0.0001 |
| Endosomal $N^1$ methyl Ψ modified mRNA vs. Non-Endosomal unmodified mRNA | *** | 0.0002 |
| Non-Endosomal unmodified mRNA vs. Non-endosomal $N^1$ methyl Ψ modified mRNA | ns | 0.9489 |
| Endosomal $N^1$ methyl Ψ modified mRNA vs. Non-endosomal $N^1$ methyl Ψ modified mRNA | *** | 0.0002 |

**Supplementary Table S4: One-way ANOVA followed by post-hoc Tukey test (of log values of nano luciferase expression) to determine statistical significance between different mRNA modifications when endosomal delivery route was chosen to deliver nano-luciferase mRNA to RAW cells. These tests support data presented in Figure 2.**

| Tukey test (post ANOVA) | Summary | Individual P Value |
|---|---|---|
| 6hrs: unmodified vs. m6A modified | *** | 0.0001 |
| 6hrs: unmodified vs. m5C modified | ns | 0.9991 |
| 6hrs: unmodified vs. pseudouridine | **** | <0.0001 |
| 6hrs: unmodified vs. $N^1$ -methyl pseudouridine | **** | <0.0001 |
| 24hrs: unmodified vs. m6A modified | * | 0.0154 |
| 24hrs: unmodified vs. m5C modified | ns | 0.9888 |
| 24hrs: unmodified vs. pseudouridine | **** | <0.0001 |
| 24hrs: unmodified vs. $N^1$ -methyl pseudouridine | **** | <0.0001 |

**Supplementary Table S5: One-way ANOVA followed by post-hoc Tukey test (of log values of nano luciferase expression) to determine statistical significance between different mRNA modifications when non-endosomal delivery route was chosen to deliver nano-luciferase mRNA to RAW cells. These tests support data presented in Figure 2.**

| Tukey test (post ANOVA) | Summary | Individual P Value |
|---|---|---|
| 6hrs: unmodified vs. m6A modified | *** | 0.0004 |

| | | |
|---|---|---|
| 6hrs: unmodified vs. m5C modified | ns | 0.9932 |
| 6hrs: unmodified vs. pseudouridine | ns | 0.8793 |
| 6hrs: unmodified vs. $N^1$ -methyl pseudouridine | ns | 0.9684 |
| 24hrs: unmodified vs. m6A modified | *** | 0.0006 |
| 24hrs: unmodified vs. m5C modified | ns | 0.9958 |
| 24hrs: unmodified vs. pseudouridine | ns | 0.8215 |
| 24hrs: unmodified vs. $N^1$ -methyl pseudouridine | ns | 0.9940 |

**Supplementary Table S6: Kruskal-Wallis test to determine if TNF-α levels are changing significantly in RAW cells following transfection with mRNA with different modifications delivered via the endosomal route and measured after 6 hours and 24 hours**

| Uncorrected Dunn's Test | Summary | Individual P Value |
|---|---|---|
| 6hrs: unmodified vs. m6A modified | ns | 0.5134 |
| 6hrs: unmodified vs. m5C modified | ns | 0.6369 |
| 6hrs: unmodified vs. pseudouridine | ns | 0.4903 |
| 6hrs: unmodified vs. $N^1$ -methyl pseudouridine | ns | 0.1882 |
| 24hrs: unmodified vs. m6A modified | ns | 0.6216 |
| 24hrs: unmodified vs. m5C modified | ns | 0.4136 |
| 24hrs: unmodified vs. pseudouridine | ns | 0.9222 |

| | | |
|---|---|---|
| 24hrs: unmodified vs. $N^1$ -methyl pseudouridine | ns | 0.3597 |

**Supplementary Table S7: Kruskal-Wallis test to determine if TNF-α levels are changing significantly in RAW cells following transfection with mRNA with different modifications delivered via the non-endosomal route and measured after 6 hours and 24 hours**

| Uncorrected Dunn's Test | Summary | Individual P Value |
|---|---|---|
| 6hrs: unmodified vs. m6A modified | ns | 0.1414 |
| 6hrs: unmodified vs. m5C modified | ns | 0.9277 |
| 6hrs: unmodified vs. pseudouridine | ns | 0.9855 |
| 6hrs: unmodified vs. $N^1$ -methyl pseudouridine | ns | 0.1783 |
| 24hrs: unmodified vs. m6A modified | ns | 0.8330 |
| 24hrs: unmodified vs. m5C modified | ns | 0.7151 |
| 24hrs: unmodified vs. pseudouridine | ns | 0.0433 |
| 24hrs: unmodified vs. $N^1$ -methyl pseudouridine | ns | 0.9207 |

**Supplementary Table S8: Kruskal-Walis test to determine if IL-6 levels are changing significantly in RAW cells following transfection with mRNA with different modifications delivered via the endosomal route and measured after 6 hours and 24 hours**

| Uncorrected Dunn's Test | Summary | Individual P Value |
|---|---|---|
| 6hrs: unmodified vs. m6A modified | ns | 0.6452 |

| | | |
|---|---|---|
| 6hrs: unmodified vs. m5C modified | ns | 0.6744 |
| 6hrs: unmodified vs. pseudouridine | ns | 0.5572 |
| 6hrs: unmodified vs. $N^1$ -methyl pseudouridine | ns | 0.9113 |
| 24hrs: unmodified vs. m6A modified | ns | 0.4740 |
| 24hrs: unmodified vs. m5C modified | ns | 0.3232 |
| 24hrs: unmodified vs. pseudouridine | ** | 0.0093 |
| 24hrs: unmodified vs. $N^1$ -methyl pseudouridine | ns | 0.6909 |

**Supplementary Table S9: Kruskal-Walis test to determine if IL-6 levels are changing significantly in RAW cells following transfection with mRNA with different modifications delivered via the non-endosomal route and measured after 6 hours and 24 hours**

| Uncorrected Dunn's Test | Summary | Individual P Value |
|---|---|---|
| 6hrs: unmodified vs. m6A modified | ns | 0.9129 |
| 6hrs: unmodified vs. m5C modified | ns | 0.7868 |
| 6hrs: unmodified vs. pseudouridine | ns | 0.5118 |
| 6hrs: unmodified vs. $N^1$ -methyl pseudouridine | * | 0.0498 |
| 24hrs: unmodified vs. m6A modified | ns | 0.7308 |
| 24hrs: unmodified vs. m5C modified | ns | 0.9116 |

| | | |
|---|---|---|
| 24hrs: unmodified vs. pseudouridine | ** | 0.0040 |
| 24hrs: unmodified vs. $N^1$ -methyl pseudouridine | ns | 0.6287 |
